# Small non-coding RNAs encapsulating mammalian cells fuel innate immunity

**DOI:** 10.1101/2025.04.07.647669

**Authors:** Xiao Jiang, Chu Xu, Enzhuo Yang, Danhua Xu, Yong Peng, Xue Han, Jingwen Si, Qixin Shao, Zhuo Liu, Qiuxiao Chen, Weizhi He, Shuang He, Yanhui Xu, Chuan He, Xinxin Huang, Lulu Hu

## Abstract

Cell surface RNAs, notably glycoRNAs, have been reported, yet the exact surface RNA compositions in different cell types remain unclear. Here, we introduce a comprehensive suite of methodologies for imaging, profiling, quantifying, and exploring the biological functions of specific surface RNAs. Utilizing these techniques, we have identified diverse non-coding RNAs present on mammalian cell surfaces. We confirm the membrane anchorage and quantify the abundance of several representative RNAs on human primary cells. Notably, we discover a significant prevalence of Y RNAs on the surfaces of human monocytes and B cells. We find that these Y RNAs on human monocyte surfaces enrich extracellular histones, regulating interleukin-6 (*IL-6*) gene expression and subsequent protein secretion upon histone stimulation via NF-κB and AP-1 activation. Our study not only presents effective approaches for investigating surface RNAs, but also uncovers a previously unrecognized immune activation pathway mediated by surface Y RNAs on monocytes.

**In brief:** A comprehensive profiling of surface RNAs across diverse mammalian blood cell types unveil abundant Y RNAs on the surface of human monocytes. Subsequent investigations uncover functional roles of surface Y RNAs on monocytes as “immune sentinel” to enrich extracellular histones, revealing a previously unknown pathway of innate immune activation.

**Highlights:** Novel methodologies for cell surface RNA mapping, validation and functional interrogation.

Sequencing of surface RNAs across various mammalian blood cell types offers detailed maps of surface RNAs.

Abundant Y RNAs localize on human monocyte surfaces and capture extracellular histones released by adjacent ruptured cells.

Surface Y RNAs enrich extracellular histones to facilitate transcription and secretion of IL-6 through NF-κB and AP-1 activation.

## Introduction

Cellular life thrives within the confinements of the cell membrane, the defining boundary that contains functional units. Traditionally recognized as a 4-nm^1^ lipid bilayer^2^ encompassing lipids, proteins, and glycans,^3-5^ the plasma membrane contains mosaic proteins embedded within the fluid lipid bilayer^6,7^ and cholesterol-enriched lipid rafts,^8^ acting as potential signalling platforms.

The ‘RNA world’ theory^9^ suggests an RNA-centric pre-cellular era, indicating RNA’s dual capacity to store genetic information and also catalyze reactions as ribozymes.^10-14^ The ‘RNA world’ theory may also suggest that RNA could be encapsulated by membrane bilayer or even be part of the membrane during the evolution of life. Advancements in imaging, sequencing, and mass spectrometry urged a re-evaluation of the plasma membrane’s composition. This prompted an inquiry into whether RNA also associates with the membrane. Indeed, a recent breakthrough discovered the presence of non-coding RNAs and fragments of protein-coding RNAs on mammalian cell surfaces.^15^ Moreover, small RNAs, including tRNAs, have been reported to be N-glycosylated at acp^3^U and displayed on cell surfaces,^16-18^ along with findings showing interactions between short guanine-rich RNAs and artificial unilamellar vesicles in *vitro*.^19^ Advances in techniques like spatial imaging assay further offered new views into glycoRNAs’ roles in cell-cell interactions during immune responses.^20^

Despite recent discoveries indicating the presence of multiple RNAs, notably glycoRNAs, residing on the plasma membrane,^15,16,18^ non-disruptive methodologies that enable comprehensive and precise profiling of global surface RNAs, particularly within rare cell populations, remain elusive. Crucial aspects, such as the potential diverse nature of surface RNAs and their interaction with the plasma membrane, necessitate thorough exploration. GlycoRNAs on the cell surface of murine neutrophils have been implicated in neutrophil recruitment to inflammatory sites and subsequent migration through endothelial cells.^21^ Additionally, RNA-binding proteins and glycoRNAs have been reported to form nanoclusters that facilitate the entry of cell-penetrating peptides.^22^ While previous studies have investigated total surface RNAs and used RNase A treatment as a control, there remains a need for effective mapping approaches and a detailed functional understanding of specific surface RNA species. To address this gap, we embarked on a comprehensive examination of surface RNAs from specific primary cell types, aiming to elucidate their connections with specific physiological functions related to the residing cell type.

We report here a set of new strategies tailored for the investigation of surface RNAs located on the outer plasma membrane across diverse cell types. We discovered abundant Y RNAs on the surfaces of human monocytes and B cells. Significantly, we focused on these Y RNAs due to their interaction with systemic lupus erythematosus (SLE) canonical autoantibodies SSA and SSB antigens Ro60 and La.^23,24^ We demonstrate the critical role of Y RNAs in facilitating innate immune activation in response to extracellular histones, a canonical damage-associated molecular pattern (DAMP).

## Results

### RNA localization on the outer membrane surface of cultured cells and live human umbilical cord blood mononuclear cells (hUCB-MNCs)

To investigate RNA localization on the outer membranes of mammalian cells, we developed an “Intact-Surface-FISH” assay, enabling surface RNA imaging while preserving cell viability and membrane integrity (Method details and Figure S1A). This method builds upon the Surface-FISH technique,^15,25^ optimizing the process by reducing the DNA probe incubation time to 30 minutes for live primary cells and cultured cell lines. The shortened incubation time is optimized to minimize endocytosis in live primary cells and cell lines during exposure to fluorescent DNA probes. For the Intact-Surface-FISH assay targeting surface RNAs, live primary cells or cell lines were used without fixation or permeabilization to preserve membrane integrity. 1 mM ATP, rather than formamide, was utilized to wash away non-specific oligo binding due to its non-denaturing properties, ensuring cell viability and allowing for subsequent confocal imaging and flow cytometry analysis with live cells.

We confirmed membrane integrity through transmission-through-dye (TTD) microscopic analysis^15,26-28^ (Method details and Figure S1B). AB9 failed to quench the fluorescence of cells stained with CellTracker Orange in both control and “Intact-Surface-FISH” treated cells (Figure S1B, upper and middle panels). However, in permeabilized cells treated with 0.2% Triton, the AB9 quencher penetrated, reducing fluorescence signals (Figure S1B, bottom panel). These results confirm that the “Intact-Surface-FISH” procedure maintains cell membrane integrity.

Incubation of the Cy3-20N (random A/T/C/G) probe with live HeLa and HEK 293T cells, revealed a clear fluorescent signal localized to the cell membrane through the Intact-Surface-FISH assay. The use of random 20N DNA probes, rather than shorter probes, was intended to ensure robust binding between complementary DNA probes and surface RNAs, thereby facilitating effective imaging. This signal was susceptible to RNase digestion (Figure S1C). Application of Intact-Surface-FISH to live human umbilical cord blood mononuclear cells (hUCB-MNCs), followed by flow cytometry analysis (Figures S1D, E) and confocal imaging (Figure S1F), further supported the presence of cell surface-anchored RNA, demonstrating the feasibility of our approach. These findings corroborate recent observations of surface RNA association with cultured cell outer membranes^15,16^ and validate our intact imaging method.

### “AMOUR” enables *in situ* amplification of outer membrane surface RNAs

Despite the successful identification of surface RNA using the high-throughput method “Surface-seq”,^15^ challenges remain in analyzing rare cell populations. Although the spatial imaging approach “ARPLA”^20^ is highly sensitive and provides detailed spatial information, it is specialized for imaging a specific type of surface RNA rather than multiple surface RNAs in a high-throughput manner. Therefore, our aim is to develop a non-disruptive technique that facilitates comprehensive and accurate profiling of global surface RNA residing on the plasma membrane, particularly focusing on rare primary cell populations. Building on promising flow cytometry and imaging observations (Figures S1C-F), we introduce a new method, termed *in-situ* <u>am</u>plification of <u>ou</u>ter membrane surface <u>R</u>NA (AMOUR), designed to profile surface RNA species while preserving cell membrane integrity. AMOUR harnesses T7 RNA polymerase’s capability to utilize both DNA and RNA as templates for *in vitro* transcription, ^29^integrating approaches we previously developed in Jump-seq.^30^

In the context of AMOUR (Figure 1A and Method details), it is crucial to maintain gentle procedures, particularly during *in-situ* amplification, to preserve cell membrane integrity. Live cultured cells or primary cells (with viability no less than 95%) were harvested without trypsin treatment. Following gentle washes with 1 × PBS, cells were pelleted by centrifuging at 500 × g at 25°C for 5 min. Then cells were fixed using 4% paraformaldehyde in PBS for 15 min at 25°C to stabilize the cell membrane. Cells were then immobilized onto magnetic ConA beads, and a double-stranded DNA probe (T7N9 oligo) containing a T7 promoter sequence with protruding nine N bases (random A/T/C/G) was annealed (Key resources table). We selected random nine-nucleotide sequences, rather than longer sequences, for annealing with surface RNAs to provide versatile binding sites that encompass a broad range of surface RNA regions, in accordance with the single-cell bisulfite sequencing strategy.^31^ Excess DNA probes were washed away using 1 × PBS containing 1 mM ATP before *in situ* amplification. T7 RNA polymerase (detergent-free), along with NTP, was introduced (final reaction condition: 0.5 × PBS and 0.5 × commercial reaction buffer to maintain acceptable osmotic pressure) to initiate *in-situ* amplification at 37 °C. The T7 RNA polymerase identifies the T7 promoter in the dsDNA probe, transcribing RNA along it, and “jumping” onto surface RNAs upon encountering the RNA-random N hybrid, thereby continuing transcription utilizing surface RNAs as templates. This process yields amplified RNA ready for subsequent reverse transcription with M-MLV Reverse Transcriptase, library construction and high-throughput sequencing (Method details). Unlike the Intact-Surface-FISH assay, which utilizes live cells for coupling with subsequent live cell flow cytometry analysis, the AMOUR method employs fixed cells. This approach is essential for working with rare primary cells, as live cells exhibit poor binding to ConA beads due to the fluidity of their plasma membranes. Fixed cells are used to ensure efficient capture for surface RNA profiling, minimizing significant cell loss caused by repetitive washing and centrifugation steps.

**Figure 1.**
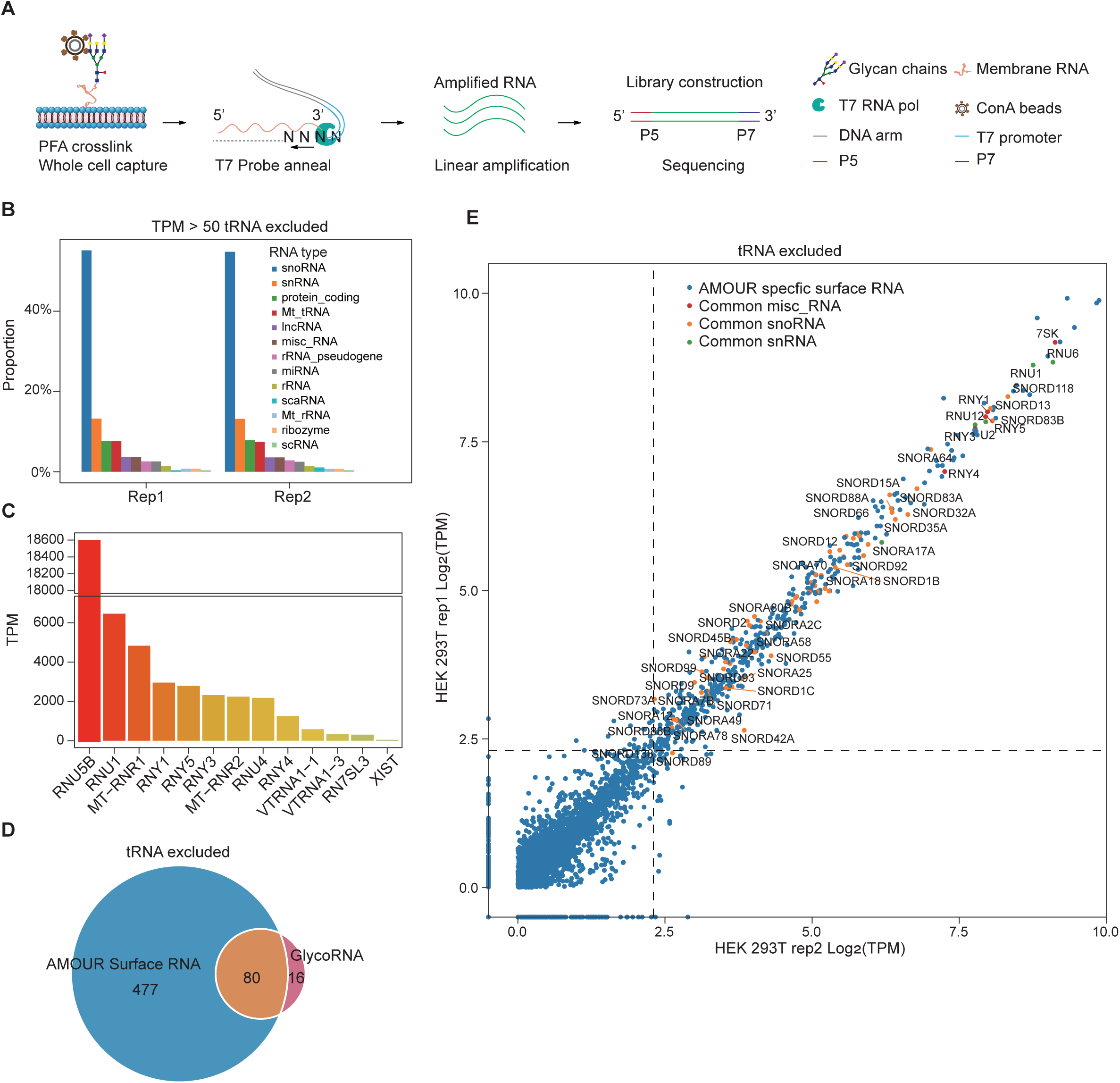
Detection of Surface RNAs via AMOUR Strategy on outer membrane of HEK 293T Cells. (A) Schematic representation detailing the workflow of the *in-situ* <u>am</u>plification of <u>ou</u>ter membrane surface <u>R</u>NA (AMOUR) technique. Fixed cells are immobilized using ConA beads, followed by hybridization with T7 probes and amplification using T7 RNA polymerase. The resulting amplified RNA undergoes library construction and subsequent sequencing. (B) Bar plot displaying the composition of RNA types present in surface RNAs of HEK 293T cells (excluding tRNA) as detected through the AMOUR method (n = 2 independent biological replicates). (C) Bar plot showcasing the transcripts per million (TPM) values of select representative surface RNA molecules identified in HEK 293T cells. (D) Venn plot demonstrating the overlap between surface RNA detected via AMOUR and glycoRNAs reported by Ryan A Flynn et al. (Cell. 2021), excluding tRNA. (E) Scatter plot highlighting the overlap between surface RNAs detected via AMOUR and glycoRNAs reported by Ryan A Flynn et al. (Cell. 2021), excluding tRNA. AMOUR-specific surface RNAs are denoted with a blue hue, while common misc_RNA, snoRNA, and snRNA are represented with red, orange, and green hues, respectively.

### Validation of AMOUR strategy

We initially examined the impact of AMOUR treatment on cell membrane integrity. Microscopic analysis following AMOUR treatment with TTD (Method details) revealed that cells retained resistance to the quencher and exhibited fluorescence intensities comparable to the control group, in contrast to cells treated with 0.2% Triton X-100, which showed significant damage (Figure S2A). Control experiments indicate that neither the Alexa Fluor 647-GAPDH antibody, Cy3-GAPDH oligo, nor the Alexa Fluor 647-T7 RNA polymerase penetrates fixed cells under AMOUR conditions (Figures S2B-D). Significant signals of Cy3 or Alexa Fluor 647 were observed only in cells permeabilized with 0.2% Triton X-100. Notably, mild permeabilization using 0.001% or 0.01% Triton X-100 resulted in minimal leakage of Alexa Fluor 647 T7 RNA polymerase, with substantial leakage occurring only following thorough permeabilization at 0.2% Triton X-100 (Figures S2D). Imaging data confirmed that complete permeabilization of fixed cells, rather than minimal damage, induces leakage of macromolecules like T7N9 oligos or T7 RNA polymerase. Since the AMOUR assay uses cells with ≥ 95% viability and preserves membrane integrity, these findings underscore the method’s practical applicability.

We then assessed the effectiveness of the AMOUR strategy for *in situ* amplification using a 325 nt biotinylated model RNA immobilized on streptavidin beads, demonstrating robust amplification efficiency and reads coverage (Figure S3A). We further applied AMOUR amplification to lysed HEK 293T cells, as well as to HEK 293T cells pre-treated with RNase A and T1 in their live states prior to fixation, to serve as additional controls. The results revealed significant differences compared to the standard HEK 293T AMOUR assay (Figure S3B). Notably, amplifying just 3% of the cell lysate (Method details) produced much stronger signals than the standard AMOUR assay, while the RNase pre-treated group exhibited a marked reduction in surface RNA signals (Figure S3C). We noted that several surface RNAs, including RNVU1 and RNU6, retained strong signals despite RNase treatment. We observed that these specific RNAs are highly structured, which protects them from RNase degradation and leads to their over-amplification by T7 RNA polymerase followed by sequencing. Transcripts enriched in the RNase-treated group displayed a greater propensity to form stable secondary structures compared to both the 3% cell lysate and AMOUR control groups (Figure S3D). Consistent with this observation, analysis of the sequence context of these enriched transcripts revealed a significant increase in G/C content (Figure S3E). Since TPM represents the relative abundance of a transcript within a population of sequenced transcripts,^32^ these highly structured surface RNAs, which are resistant to RNase, exhibit strong signals during sequencing.

Transcripts identified by AMOUR with a TPM (transcripts per kilobase per million mapped reads) value greater than 50 are considered positive hits. TPM values for snoRNA, scaRNA, rRNA, and ribozyme in the AMOUR control group are significantly lower than those in the 3% cell lysate group, while mitochondrial tRNAs are significantly enriched in the AMOUR group (Figure S3F). The TPM values of representative small RNAs are higher than those of protein-coding RNAs identified by AMOUR (Figure S3G). Mitochondrial tRNAs, including MT-TA, MT-TN, and small RNAs such as RNY1, RNY4, and VTRNA1-1, are predominantly enriched in the AMOUR control group (Figure S3G, left panel). In contrast, RNY3, RNY5, and all representative protein-coding RNAs—such as H3C1, H4C3, RANBP1, RPL13, RPL36A, RPL41, and TMEM107—are enriched in the cell lysate group (Figure S3G, right panel). The TPM values of all representative RNAs exhibit a marked reduction in the RNase pre-treated group, followed by AMOUR. Notably, mitochondrial tRNAs retain substantial TPM values in the RNase pre-treated group, likely due to their stable secondary structures (Figure S3G, left panel). Quantitative analysis further reveals that the surface RNAs identified by AMOUR exhibit a distinct pattern, rather than representing leakage from the intracellular compartment. The prevalent presence of surface tRNAs detected aligns well with previous reports indicating that tRNAs are highly glycosylated and displayed on the cell surface.^16,18^ Previous research has shown that human tRNA^Sec^ associates with HeLa membranes, cell lipid liposomes, and synthetic lipid bilayers,^33^ consistent with our findings. This supports the need for further investigation into the role of surface tRNAs.

### Surface RNAs on the outer membrane of HEK 293T and Hela Cells

Surface RNAs from the outer membrane of HEK 293T cells were profiled using the AMOUR strategy (Table S1), revealing consistent patterns across two biological replicates (Figure S4A). This consistency underscores the reliability and stability of our approach. Notably, the majority of surface RNAs tethered to the outer membrane consist of tRNAs and other non-coding RNAs, with protein-coding RNAs forming a minority (Table S1 and Figure S4B). To investigate the diversity of surface RNAs, we excluded tRNAs due to their limited species diversity and focused the subsequent data analysis on non-tRNA surface RNAs. It turns out that non-tRNA surface RNAs encompass diverse categories, including small nucleolar RNAs (snoRNA) such as C/D box and H/ACA box snoRNAs, as well as small nuclear RNAs (snRNA) covering key spliceosome components like U5, U6, U1, U2, U7, and U11, all of which have previously been reported as glycosylated.^16^ Additionally, various other non-coding RNAs including mitochondria tRNA (mt-tRNA), long non-coding RNA (lncRNA), miscellaneous small RNA (misc-RNA), rRNA-pseudogene, microRNA (miRNA), rRNA, small Cajal body-specific RNA (scaRNA), mitochondria rRNA (mt-rRNA), ribozyme, and small cytoplasmic RNA (scRNA) are also enriched (Figures 1B and S4C).

Representative examples illustrating different types of surface RNAs (Figures 1C and S4D) suggest potential diverse roles. Among these, RN7SL3 RNA stands out as a key component of the signal recognition particle (SRP), responsible for the transport of membrane proteins to the plasma membrane, an association consistent with its surface localization.^34^ Furthermore, the interaction of VTRNA1-1 with P62 and its implicated role in autophagy inhibition prompt compelling inquiries into its function when localized on the cell surface.^35^ Additionally, the presence of mitochondria rRNA MTRNR1/2 and the lncRNA XIST, known for initiating X chromosome silencing, is surprising. We were particularly intrigued by several snRNAs, including U1/2/4/5/6, and Y family RNAs, given the significant role of spliceosomal RNAs in constituting the Sm antigen^36^ and the binding of Y RNAs to SLE canonical autoantibodies SSA and SSB antigens Ro60 and La,^23,24^ respectively. Surface RNAs identified through the AMOUR strategy encompass the majority of previously reported glycoRNAs^16^ (Figures 1 D, E and S4E, F).

The surface RNA profile of HeLa cells closely resembles that of HEK 293T cells (Figure S4G), although HeLa cells exhibit a higher proportion of tRNAs and a lower proportion of snoRNAs on their surface compared to HEK 293T cells (Table S1 and Figures S4B, H). Quantitative analysis shows that the TPM values of HEK 293T surface snoRNA, scaRNA, and miRNA are higher than that of HeLa surface RNAs, while surface mitochondria tRNAs tend to be more abundant in HeLa cells (Figure S4I).

### Validation of ARMOUR results

To ensure both sensitivity and specificity, we developed two complementary approaches for surface RNA profiling in conjunction with ARMOUR. The first approach utilizes a wheat germ agglutinin (WGA) pull-down assay (WGA-Pd, Method details and Figure S5A), in which Biotin-WGA is used to capture plasma membranes enriched in N-acetylglucosamine and N-acetylneuraminic acid (sialic acid). This is followed by surface RNA extraction and reverse transcription using M-MLV Reverse Transcriptase. We define surface RNAs as any RNA molecules displayed on the outer cellular membrane, whether glycosylated or not, and regardless of whether they directly or indirectly anchor to the cell surface. Given the widespread application of wheat germ agglutinin (WGA) as a cell membrane marker and its established efficacy in isolating glycosylated RNAs,^37^ we utilized Biotin-WGA to selectively pull down all RNAs associated with the cell membrane via glycosylated proteins, glycans, or glycosylated lipids, thereby serving as a comprehensive positive control. A comparison of enriched surface RNAs obtained through WGA and ARMOUR reveals a notable consistency (Figures S5C, D), supporting the specificity of AMOUR. Considering that we utilized 1 million cells for AMOUR compared to 20 million cells for WGA-Pd during surface RNA identification (Method details), AMOUR demonstrates superior sensitivity over WGA-Pd when working with limited primary cell samples. IGV tracks displaying the reads coverage of RNY1/3/4/5 identified by the three approaches further demonstrate that AMOUR and WGA-Pd generate complete coverage (Figure S5F). Leveraging T7 RNA polymerase’s *in-situ* linear amplification and strand displacement capabilities, AMOUR proves to be a highly sensitive method for profiling surface RNA, especially when working with limited primary cell numbers. Notably, the sensitivity and specificity of AMOUR, without disrupting the cell membrane, allows us to map surface RNA landscapes across diverse primary cell types.

### Verification of identified surface RNA molecules with cultured Hela cells and hUCB-MNCs

Reports have indicated the presence of surface RNAs,^15^ notably glycoRNAs^17,18^ on the plasma membrane. Current imaging studies have confirmed the localization of specific surface RNAs on both cultured cell lines,^15,16,20,38^ and peripheral blood mononuclear cells (PBMCs).^15^ To validate the membrane localization of previously identified surface RNAs^16^ and those identified via AMOUR, we employed the self-developed Intact-Surface-FISH assay (Method details and Figure S1A). Confocal imaging data clearly demonstrate the localization of small RNA RNY5 and mt-rRNA MT-RNR1/2 on the surface of cultured HeLa cells (Figure 2A), labeled with complementary Cy3 probes (Key resources table). We designed the oligonucleotides to be 35 to 40 nucleotides in length, with a GC content ranging from 40% to 60%. Sequences containing continuous CG trinucleotides were excluded. For small RNA targets, one to two oligonucleotide sequences were designed to cover the entire transcript. For longer transcripts, two to three oligonucleotide sequences were designed to cover two to three exons (Key resources table). Using flow cytometry, we quantified the proportion of live hUCB-MNCs displaying several highlighted surface RNAs, including small RNA RNY5, mitochondrial rRNAs MT-RNR1/2, spliceosomal RNA U5, and lncRNA XIST (Figure 2B). These representative surface RNAs, which encompass various RNA types, were selected due to their documented essential biological functions. We further confirmed the colocalization of these surface RNAs with the plasma membrane in primary hUCB-MNCs (Figure 2C). Notably, Cy3-probes complementary to lambda DNA and GAPDH showed no signal, further reinforcing the assay’s specificity. Our flow cytometry analysis and imaging results unequivocally confirm the membranal localization of Y family RNA Y5, spliceosomal snRNA U5, mt-rRNAs, and lncRNA XIST. Notably, primary hUCB-MNCs exhibit a broader variety of surface RNA types compared to cultured HeLa cells (Figures 2A-C), potentially due to their greater cellular diversity.

**Figure 2.**
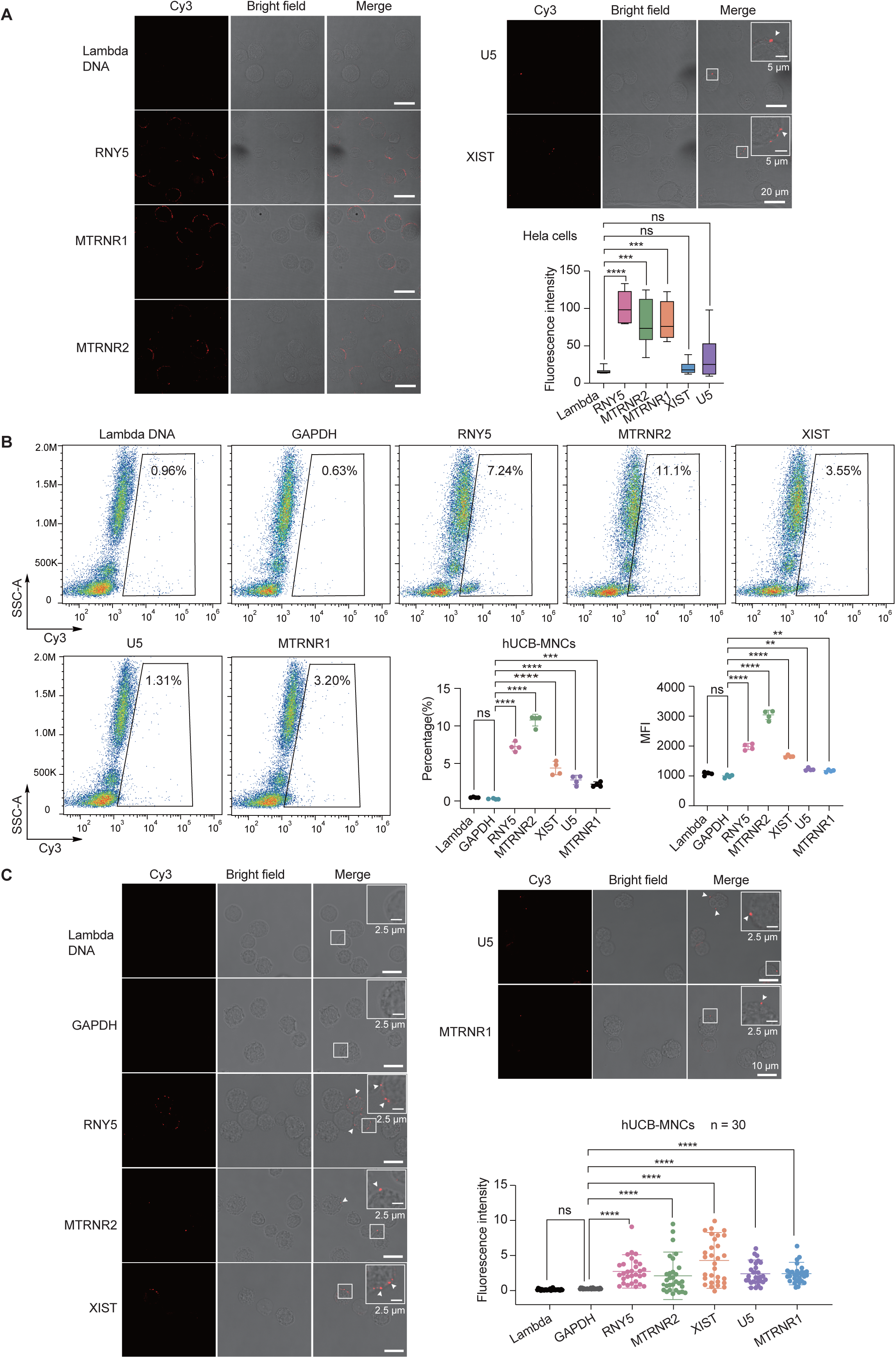
Visualization of surface RNA molecules anchored to the outer membrane surface of mammalian cells. (A) Confocal imaging of HeLa cells stained with Cy3-labeled oligonucleotides complementary to specific surface RNAs using Intact-Surface-FISH. White arrowheads denote the precise locations of Intact-Surface-FISH signals. Quantitative analysis of fluorescence intensity for each Cy3-DNA probe labeling is provided. (B) Fluorescence labeling of live hUCB-MNCs with Cy3-Lambda DNA and GAPDH controls, as well as Cy3-DNA probes complementary to RNA Y5, MTRNR2, XIST, U5, and MTRNR1, using Intact-Surface-FISH. The gated region highlights the cell population exhibiting specific RNAs on the cell surface. Quantitative analysis of cell fraction percentages and mean fluorescence intensity within the Cy3-positive region is provided for each Cy3-DNA probe labeling. (C) Confocal imaging of hUCB-MNCs stained with Cy3-labeled oligonucleotides complementary to representative surface RNAs using Intact-Surface-FISH. White arrowheads indicate the specific locations of Intact-Surface-FISH signals. Quantitative analysis of fluorescence intensity per cell for each Cy3-DNA probe labeling is provided. All flow cytometry analyses represent four independent experiments, and confocal imaging data represent three independent experiments with similar results, shown as mean ± s.d., and analyzed using an unpaired two-tailed Student’s *t*-test; ns: not significant, **p < 0.01, ***p < 0.001, ****p < 0.0001.

To further validate membrane integrity during the Intact-Surface-FISH assay, we co-incubated Cy3-Y5 and Cy5-GAPDH and performed Intact-Surface-FISH on both live and fixed, permeabilized hUCB-MNCs. Flow cytometry (Figure S6A) and confocal imaging (Figure S6B) revealed that small RNA Y5, rather than mRNA GAPDH, is prominently displayed on the surface of live hUCB-MNCs. The surface fluorescence signals of these representative RNAs distinctly differ from the intracellular signals observed after fixation and permeabilization utilizing conventional RNA-FISH. (Figures S6A-C). Please note that for imaging RNAs localized within the nucleus after fixation and permeabilization, a prolonged incubation period of no less than 6 hours at 37°C is essential to ensure thorough hybridization of Cy3-DNA probes with nuclear RNAs (Method Details). The negligible fluorescent signal of Lambda DNA in both intact and permeabilized cells confirms the specificity of the imaging (Figures 2A-C and S6C). The signal of U5 RNA, whether using Intact-Surface-FISH (Figures 2A-C) or conventional FISH (Figure S6C), is lower compared to other detected RNAs due to its highly folded structures.

Our validation of representative surface RNAs highlights the practicality of the AMOUR strategy. Notably, no current approach—whether *in situ* amplification using intact cells like AMOUR, pull-down with Biotin-WGA, or biotinylation of all surface RNA molecules followed by pull-down^16^—can fully eliminate the risk of intracellular RNA contamination. Instead, all existing methods strive to enrich potential surface RNA candidates as accurately as possible. AMOUR offers a practical tool for identifying previously uncharacterized, abundantly enriched surface RNA species in limited primary cell populations, while Intact-Surface-FISH enables the visualization and quantification of specific surface RNA molecules on live cell membranes across diverse biological contexts. The integration of AMOUR with Intact-Surface-FISH enables comprehensive screening, improves accuracy, supports statistical analysis, and establishes a foundation for subsequent functional studies of surface RNAs in primary cells.

### Intracellular RNAs are transported and presented on the cell Surface

Supplementation with Brefeldin A, a known inhibitor of membrane trafficking and exocytosis, significantly reduced the surface RNA signal on Raji cells— which are known for their robust secretion system due to their B cell origin—as demonstrated by both flow cytometry (Figure S7A) and confocal microscopy (Figure S7B). These findings suggest that at least a portion of surface RNA may be transported to the cell membrane via vesicle trafficking. Further studies are required to solidify this hypothesis.

### Comprehensive mapping of surface RNAs residing on plasma membrane of mammalian blood cells

To explore the functional significance of surface RNAs, we utilized the AMOUR approach to characterize these molecules present on the outer membrane of *Homo sapiens* and *Mus musculus* blood cells. To investigate the general characteristics and abundance of surface RNAs on human blood cells, we utilized neonate hUCB-MNCs rather than adult PBMCs (peripheral blood mononuclear cells), due to the higher abundance of hematopoietic stem and progenitor cells (HSPCs) in neonate hUCB-MNCs and their significantly lower biological variability compared to PBMCs isolated from adult donors. Our investigation encompassed various blood cell types, including B cells, T cells, HSPCs, Natural Killer (NK) cells, and monocytes. Our approach successfully delineated a detailed landscape of surface RNAs across mammalian blood cells (Tables S2-5). While we excluded the conserved surface tRNAs, the investigation of the most abundant non-tRNA surface RNAs (top 150) revealed a predominance of non-coding RNAs in both human (Figure S8A) and mouse species (Figure S8B). Correlation analysis demonstrates excellent consistency among the three biological replicates, with Pearson correlation coefficients ranging from 0.84 to 0.99 (Figures S9, S10). Notably, in the non-tRNA group, snRNA, snoRNA, rRNA, and mt-rRNA made up the largest proportions, with misc-RNA and miRNA also showing substantial representation. Interestingly, ribozyme RNAs were detected on the surface of both human and mouse HSPCs (Figures S8A, B), suggesting their potential biological relevance. Additionally, surface RNA types and their abundance across different cell types within the same species displayed a high degree of similarity (Figure S8C).

Our further analysis focused on relatively abundant non-tRNA surface RNAs in each cell type of both human (Figures 3A, B and Table S2) and mouse (Figures 3C, D and Table S3). Notably, immune cell types exhibited enrichment in inflammation, innate, and adaptive immune-related functions (Figures 3B, D), whereas human HSPCs highlighted miRNA and translation-related functions (Figure 3B) due to the presence of abundant miRNA and snoRNA on their membrane surfaces. The cell type-specific surface RNAs provided by our study form a crucial basis for further investigations into biological functions. Representative highly abundant surface RNAs, spanning spliceosomal U RNA, ribosomal RNA, microRNA, mitochondria tRNA, and protein coding RNA, were meticulously identified for each distinct human (Figure S8D) and murine (Figure S8E) blood cell type. The abundant presence of surface RNAs on human monocytes is consistent with previous report.^15^

**Figure 3.**
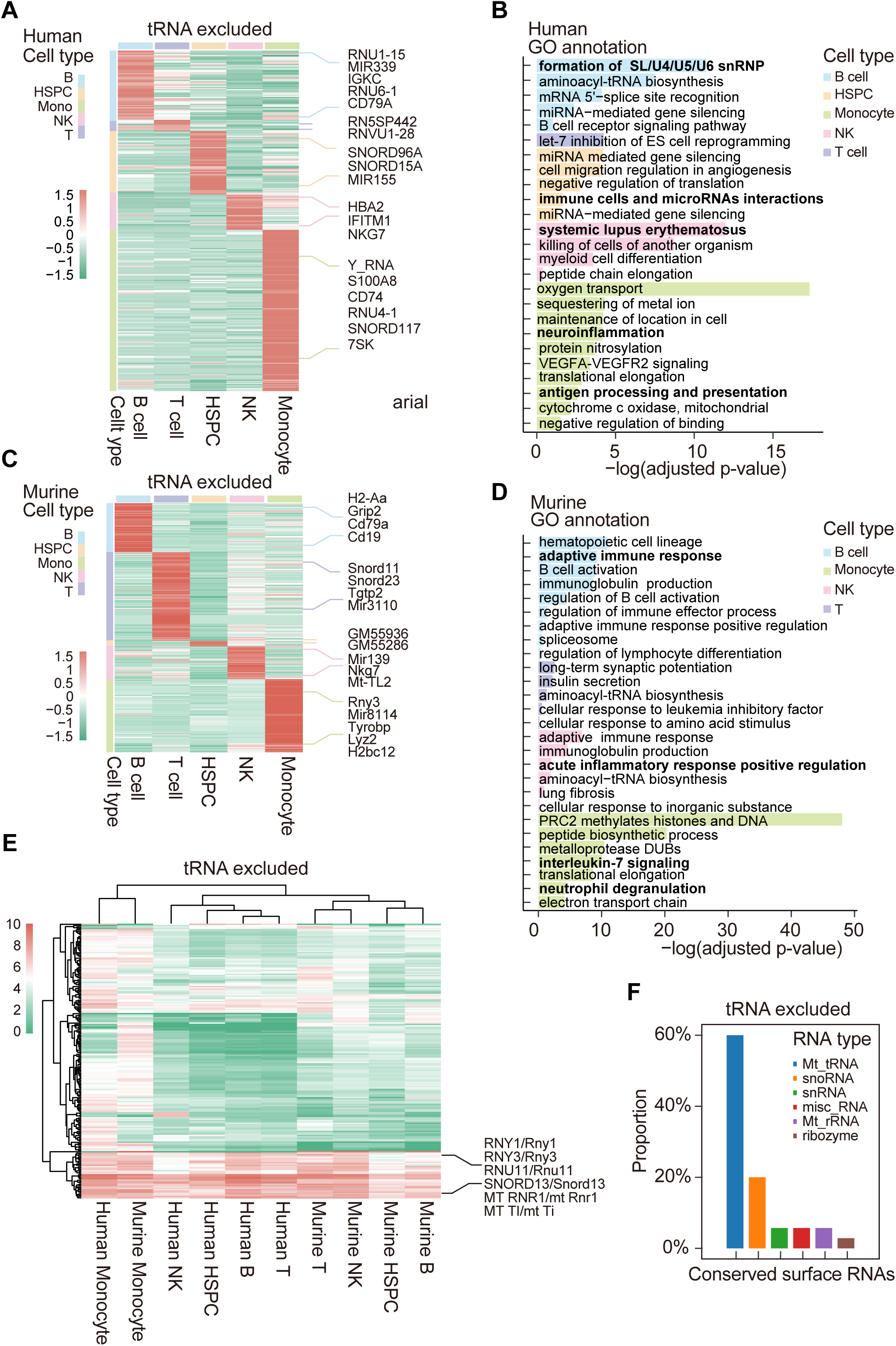
Surface RNA profiles across diverse human and murine blood cell types. (A) Heatmap illustrating distinctive surface RNA clusters characterizing various human blood cell types. Each row represents a specific cell type, and columns correspond to individual RNA clusters. Highlighted representative RNA signatures elucidate unique expression patterns across these cell types (excluding tRNA). (B) Gene Ontology (GO) analysis revealing enriched gene functions associated with observed surface RNAs across diverse human blood cell types (excluding tRNA). (C) Heatmap displaying specific surface RNA clusters identified within distinct murine blood cell subsets (excluding tRNA). Highlighted representative RNA signatures emphasize discernible expression patterns within these subsets. (D) GO analysis delineating enriched gene functions linked to surface RNAs identified across diverse murine blood cell types (excluding tRNA). (E) Heatmap spotlighting highly abundant surface RNA clusters conserved across all analysed blood cell types in both human and mouse (excluding tRNA). Highlighted representative RNA signatures underscore shared expression patterns among these cell types. (F) Bar plot illustrating the RNA types of conserved surface RNAs across all analyzed blood cell types in both human and mouse (excluding tRNA).

Remarkably, several non-tRNA surface RNA molecules exhibited high expression patterns across all blood cell types in both human and mouse (Figures 3E, S8F and Table S4), indicating evolutionarily conserved biological functions. These “housekeeping” surface RNAs included Mt-tRNA, snoRNA, snRNA, misc-RNA, Mt-rRNA, and ribozyme (Figure 3F), with selected representatives highlighted (Figure S8F).

We then focused on surface tRNAs that remained conserved across cell types and between human and murine species. The analysis revealed that surface tRNA expression is abundant in all blood cell types (Table S5). We delineated cell-type-specific surface tRNAs in humans (Figure S8G) and mice (Figure S8H), as well as those conserved across various cell types and species (Figure S8I). Notably, both human and mouse HSPCs exhibit a significantly broader spectrum of tRNA types (Figures S8G, H) compared to other cell types, suggesting the putative specialized functional roles of surface tRNAs within the context of HSPCs.

Human UCB T cells exhibit fewer surface RNAs compared to B cells and monocytes (Figure 3A and Figure S8G). Total RNA sequencing reveals that the TPM values of transcripts across various RNA categories in T cells are comparable to those in B cells and monocytes (Figure S8J). This observation suggests that the reduced abundance of surface RNAs in T cells is not attributable to differential gene expression. We hypothesize the presence of specific, yet unidentified, surface transport mechanisms.

### Y RNAs are enriched on the surface of human monocytes and B cells

The identification of Y RNAs on the surfaces of immune cells (Figure S8F) prompted an investigation into the specific cell types harboring these surface-bound Y RNAs. Using the Intact-Surface-FISH strategy, we incubated human PBMCs with Cy3-DNA probes designed to target a Y RNA sequence mix (Y1/3/4/5), with Cy3-DNA probes complementary to GAPDH serving as a negative control (Method details). Flow cytometry analysis revealed the presence of Y RNAs on the surface of monocytes and B cells in human PBMCs (Figures 4A, B). In contrast, Cy3-GAPDH DNA, used as a negative control, showed negligible signal. The fluorescence intensity of Y RNAs was higher on the surface of monocytes than on B cells, highlighting their potential biological role in innate immunity. Confocal imaging results confirmed the surface anchorage of Y RNAs on human PBMCs (Figure 4C).

**Figure 4.**
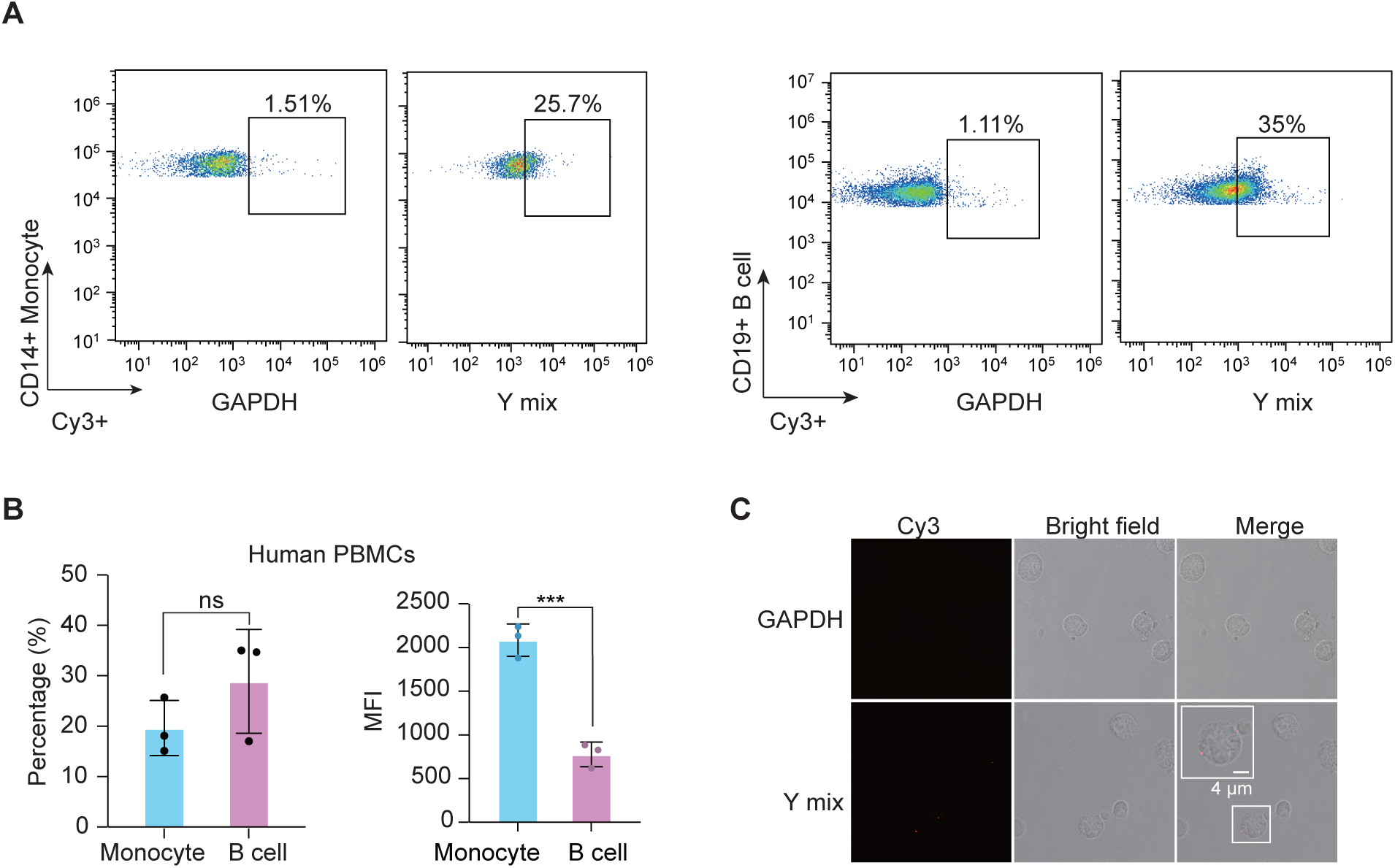
Enrichment of surface Y RNAs on the membranes of human monocytes and B cells. (A) Flow cytometry analysis of human PBMCs stained with Cy3-DNA probes complementary to surface Y RNAs utilizing Intact-Surface-FISH, followed by antibody staining for the monocyte marker CD14 (left panel) and B cell marker CD19 (right panel). A Cy3-DNA probe complementary to GAPDH was used as a control. (B) Quantitative analysis of the percentage of cell fraction (left panel) and mean fluorescence intensity (right panel) of human monocytes and B cells displaying surface Y RNAs. (C) Confocal imaging data demonstrating the membrane anchorage of surface Y RNAs on human PBMCs. Flow cytometry analysis and imaging experiments were conducted independently in triplicate, consistently yielding reproducible results across all repetitions. Data are presented as the mean ± standard deviation (s.d.) and were analyzed using an unpaired two-tailed Student’s t-test; ns: not significant, ***: p < 0.001.

For method validation and to illustrate the general surface RNA landscape, we selected hUCB-MNCs due to their uniformity across different neonate donors. Conversely, we chose PBMCs over hUCB-MNCs to investigate the abundance of surface Y RNA in specific cell types from adult donors, as PBMCs contain mature immune cells, making them more suitable for subsequent functional studies.

### Y RNAs interact with extracellular histones

Intrigued by the potential functions of surface RNAs on monocytes, we developed the <u>S</u>urface <u>R</u>NA Interacting protein <u>C</u>apture (SRIC) assay. This assay was adapted from the RIC^39^ and eRIC^40^ techniques for RNA interactome profiling (Figure S11A and Method details). Live cells were incubated with biotinylated LNA-modified DNA probes at 37°C for 30 minutes, followed by washing to remove excess DNA probes. Cells with biotin-LNA modified DNA probes annealed to surface RNAs were subjected to UV irradiation at 254 nm, while the control group was not irradiated. The brief 30-minute incubation aimed to minimize endocytosis, and the UV irradiation was used to covalently crosslink the RNA-protein complexes. Subsequently, cells were lysed, and surface RNA-binding proteins (RBPs) were isolated using streptavidin beads for peptide fingerprint analysis. Proteins with a fold change (mass spectrometry area (+UV/-UV)) ≥ 1.5 were considered potential hits, ensuring that only surface RBPs directly interacting with surface RNA were identified.

Peptide fingerprint analyses, based on two biological replicates, revealed that Y RNAs interact with histones, as well as with RNA binding proteins NCL, WDR43, and PTBP1 (Figures S11B and Table S6). The identification of histones is particularly noteworthy given their well-established roles as autoantigens in autoimmune diseases.^41^ *In vitro* studies demonstrated that Y5 RNA exhibits binding affinity to H2A-H2B and H3-H4 histones, with dissociation constants (Kd) of 232.2 nM and 51.56 nM, respectively (Figure S11C).

### Surface Y RNAs facilitates innate immune activation induced by extracellular histones

Canonical histones play crucial roles in organizing DNA into chromatin and serve as molecular switches. Their presence in extracellular milieu during diseases associated with cellular damage suggests their role as Damage Associated Molecular Pattern (DAMP) signals upon release from damaged cells or neutrophils through neutrophil extracellular trap formation (NETosis).^42,43^ Extracellular histones have been identified in extracellular fluids, where they influence innate immunity and cytokine levels via TLR2 and TLR4 pathways.^44-47^ Given the essential role of Toll-like receptors in the innate immune response and the prevalent presence of Y RNAs on the surfaces of human monocytes (Figures 4A-C), we hypothesize that surface-anchored Y RNAs on monocytes may play a critical role in the innate immune response to histone stimulation.

To validate this hypothesis, we evaluated the impact of surface RNA depletion on the differentiation of THP-1 monocytes. We optimized the experimental conditions for total surface RNA depletion using RNase A and T1, and further developed targeted Y RNA degradation using RNase H (Figure S12A), which we name RNase <u>H</u>-<u>A</u>ssisted <u>S</u>urface RNA <u>T</u>argeted <u>E</u>limination, or “HASTE.” Live THP-1 monocytes were incubated with DNA probes complementary to specific surface RNAs, and then treated with 0.1 U μL^-1^ RNase H at 37 ℃ for 30 minutes. Confocal imaging confirmed that HASTE effectively eliminated surface Y RNAs on THP-1 monocyte surfaces (Figure S12B). Flow cytometry analysis coupled with Intact-Surface-FISH revealed that HASTE treatment resulted in approximately a 50% reduction of Y RNAs on the surface of hUCB-MNCs (Figure S12C). Furthermore, confocal imaging in combination with Intact-Surface-FISH demonstrated a significant decrease in fluorescent signals of Y RNAs on the hUCB-MNCs surface post-HASTE treatment (Figure S12D). Further experiments demonstrated that surface RNA depletion did not significantly impact THP-1 cell number, size, viability, or differentiation into macrophages (Figures S13A-D). Depletion of total surface RNA or targeted elimination of Y RNA did not induce IL-6 or IL-10 gene expression (Figure S13E). Our results underscore the non-disruptive nature of surface RNA treatment.

We then investigated the impact of surface RNAs, particularly Y RNAs, on extracellular histone capture. We incubated freshly isolated human PBMCs with Alexa Fluor 647-labeled calf thymus histones at a concentration of 2 μg/mL for 30 minutes to validate the interaction between extracellular histones and human PBMCs, and to assess the dependency on surface Y RNAs. Flow cytometry analysis shows that, after a 30-minute incubation, extracellular histones were predominantly captured by monocytes (Figure 5A). Depletion of total surface RNAs or targeted degradation of surface Y RNAs reduced the ratio and fluorescence intensity of monocyte binding to extracellular histones by approximately 50% (Figure 5A). In contrast, B cells and other cell types in human PBMCs displayed minimal surface RNA-dependent Alexa Fluor 647 histone enrichment compared to monocytes (Figures 5B, C). Confocal imaging data confirmed the capture of Alexa Fluor 647-labeled histone by live human PBMCs (Figure 5D). Given that histones bind DNA without sequence preference, we speculate that other surface RNAs may also interact with extracellular histones. Due to the prevalent abundance of Y RNAs on the cell surface, human PBMC monocytes exhibit a significant dependence on surface Y RNAs for capturing extracellular histones.

**Figure 5.**
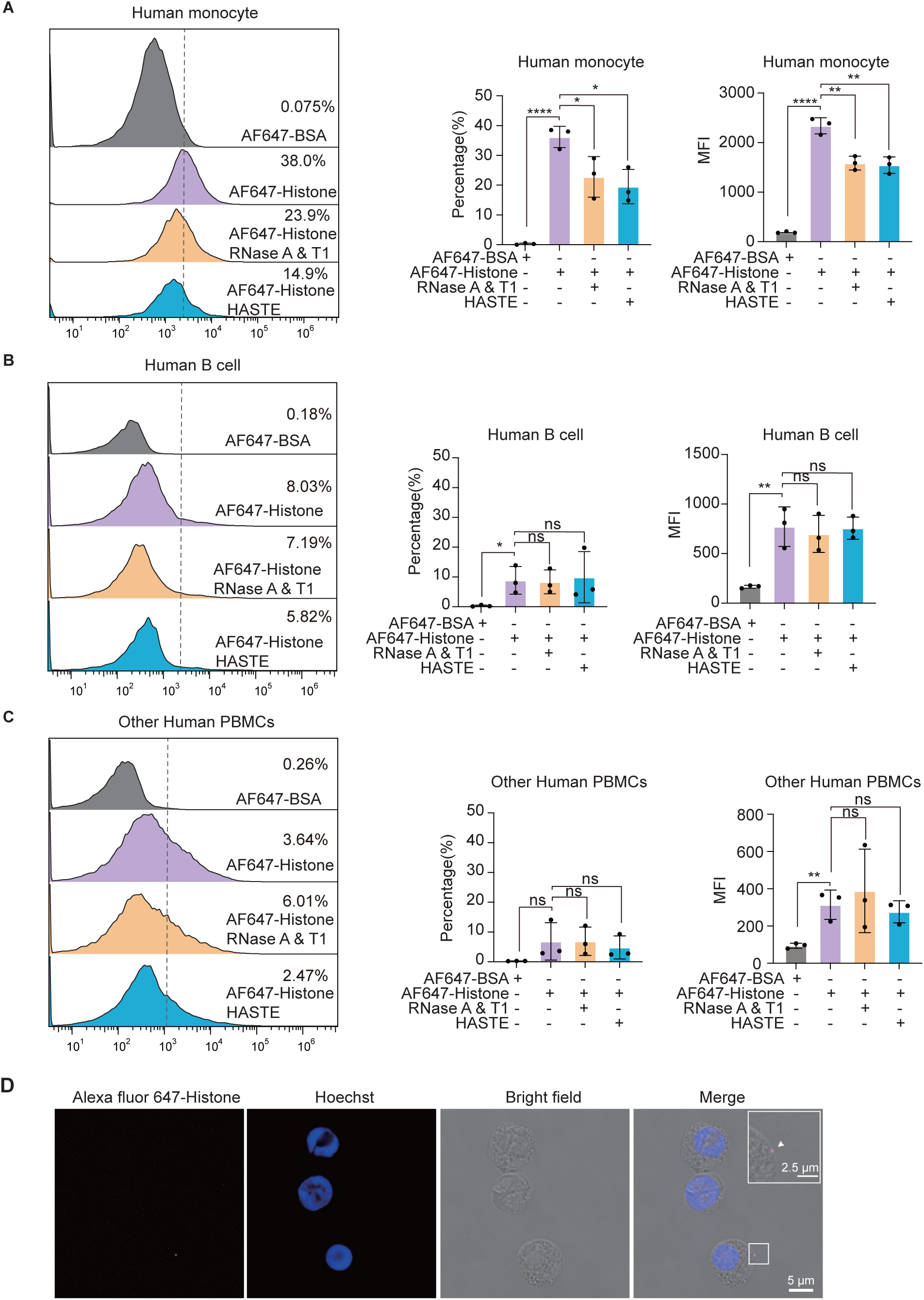
Surface Y RNAs-dependent enrichment of extracellular histones in monocytes of human PBMCs, but not B Cells or other cell types. (A-C) Fluorescence-activated cell sorting (FACS) analysis of untreated human PBMCs, as well as human PBMCs pre-treated with RNase A & T1 or HASTE utilizing RNase H and DNA probes complementary to surface Y RNAs (Y1/3/4/5), followed by incubation with Alexa Fluor-647-labeled calf thymus histones and subsequent antibody staining for the monocyte marker CD14 (A) and B cell marker CD19 (B). Alexa Fluor-647-labeled BSA was used as a control. The left panel shows the FACS results, with the dashed vertical line indicating the Alexa Fluor-647-high population. Quantitative analysis of the percentage of cell fractions (middle panel) and mean fluorescence intensity (right panel) of human monocytes (A), B cells (B) and non-B or monocyte PBMC cell types (C) within the Alexa Fluor 647-high region is shown for each treatment condition. (D) Confocal imaging of human PBMCs incubated with Alexa Fluor 647-labeled histones. White arrowhead indicates the specific location of the Alexa Fluor 647-histone signal. All imaging and flow cytometry experiments were conducted independently in triplicate, with similar results observed across all repetitions. Data are presented as the mean ± standard deviation (s.d.) from three independent experiments and were analyzed using an unpaired two-tailed Student’s t-test; ns: not significant; *: p < 0.05; **: p < 0.01, ****: p < 0.0001.

We next investigated the impact of surface RNAs, particularly Y RNAs, on cytokine production in response to histone supplementation. Short exposure (6 h) to histones resulted in elevated cytokine expression, notably IL-6. We observed that both total surface RNA and targeted surface Y RNA depletion (Figure 6A), as well as the TLR2/4 inhibitor Sparstolonin B (Figure 6B), significantly reduced *IL-6* gene expression. The suppressive effect of Sparstolonin B on histone supplementation (Figure 6B) reinforces the concept that TLR2/4 act as the principal receptors governing the cellular response to extracellular histones, consistent with previous research findings.^47-49^ Validation at the protein level via ELISA assays revealed that targeted Y RNA degradation for 30 minutes significantly attenuated IL-6 secretion, maintaining suppression for up to 72 hours (Figure 6C).

**Figure 6.**
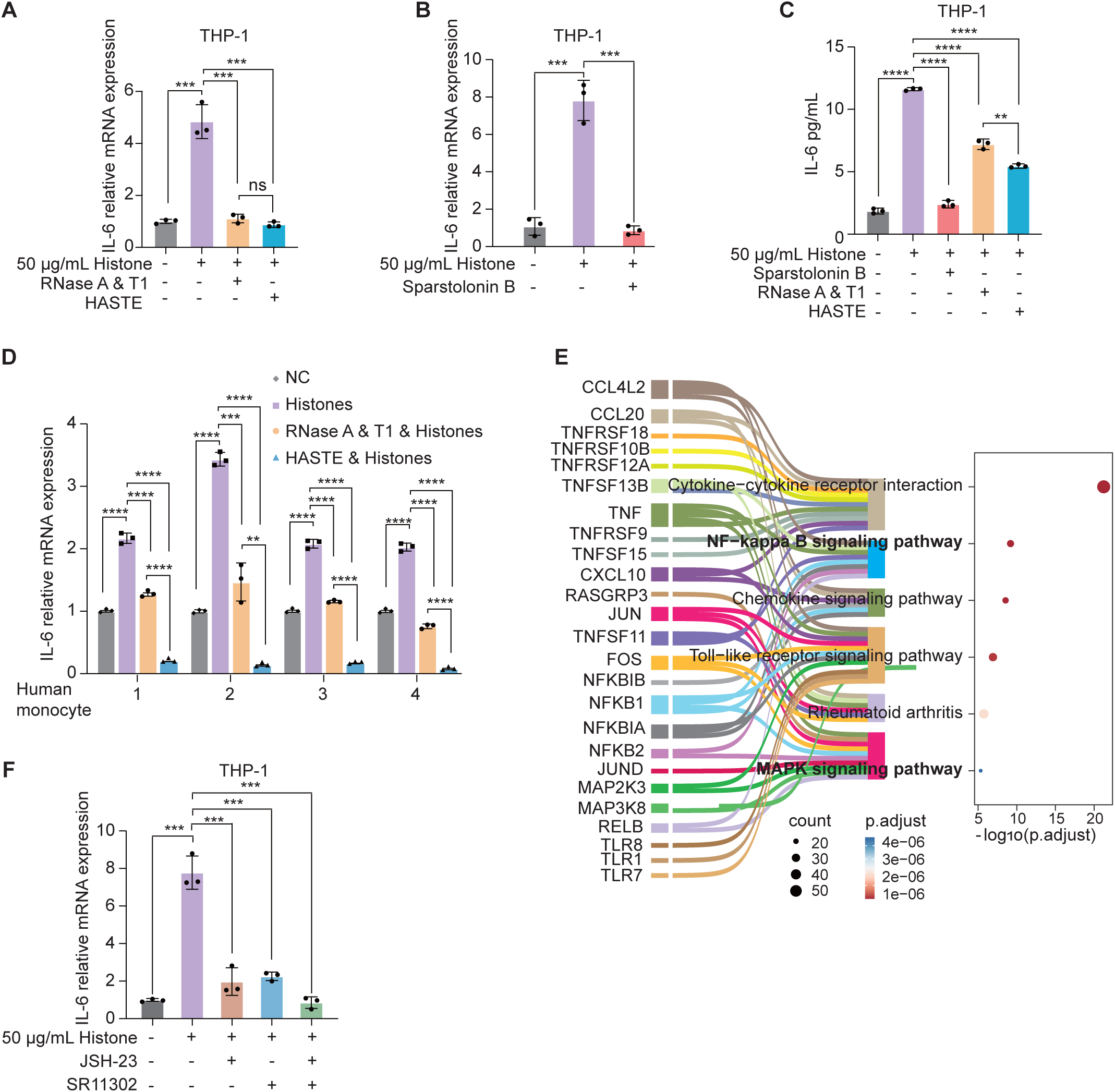
Surface Y RNAs on human monocytes promote IL-6 transcription and secretion. (A) mRNA levels of IL-6 in THP-1 macrophages cultured in FBS-free medium after 6 hours of supplementation with 50 μg mL^-1^ calf thymus histones, following various pre-treatments, as determined by RT-qPCR. Data normalized to the untreated control group, n = 3. (B) mRNA levels of IL-6 in THP-1 macrophages cultured in FBS-free medium following 6 hours of supplementation with 50 μg mL^-1^ calf thymus histones, with or without the addition of TLR2/4 inhibitor Sparstolonin B, as determined by RT-qPCR. Data normalized to the untreated control group, n = 3. (C) Protein levels of IL-6 secreted in the supernatant after 72 hours of supplementation with 50 μg mL^-1^ calf thymus histones, after various pre-treatments, as determined by ELISA. Data normalized to the untreated control group, n = 3. (D) mRNA levels of IL-6 in human monocytes purified from four human PBMC samples after 6 hours of supplementation with 50 μg mL^-1^ calf thymus histones, following various pre-treatments, as determined by RT-qPCR. NC: untreated control group; PC: human monocytes treated with calf thymus histones without pre-treatments; RNase A & T1: human monocytes treated with RNase A and T1 for 30 minutes prior to calf thymus histones supplementation; HASTE: human monocytes treated with RNase H and Y RNA complementary DNA probes for 30 minutes prior to calf thymus histones supplementation. Data normalized to the NC group, three technical replicates. (E) Sanky-bubble plot illustrating the results of KEGG (Kyoto Encyclopedia of Genes and Genomes) enrichment analysis, elucidating the critical biological pathways activated in response to histone stimulation in THP-1 macrophages. (F) mRNA levels of IL-6 in THP-1 macrophages cultured in FBS-free medium after 6 hours of supplementation with 50 μg mL^-1^ calf thymus histones, with or without the addition of NF-κB inhibitor JSH-23 or AP-1 (activator protein 1) inhibitor SR 11302, as determined by RT-qPCR. Data normalized to the untreated control group, n = 3. All data are shown as mean ± s.d. and were analysed by an unpaired two-tailed Student’s *t*-test; ns: not significant; *: p < 0.05; **: p < 0.01; ***: p < 0.001; ****: p <0.0001.

To further elucidate the physiological role of surface Y RNAs in primary cells, we evaluated the efficacy of surface RNA depletion in human monocytes isolated from PBMC samples upon 6-hour histone supplementation. The inhibitory effects observed with total surface RNA depletion and targeted Y RNA elimination in human monocytes paralleled those seen in THP-1 macrophages, with Y RNA degradation resulting in significantly greater suppression of *IL-6* and *IL-10* expression (Figures 6D and S14A). The mRNA levels of *IL-6* and *IL-10* in the Y RNA depletion group were even lower than those in the control group without histone treatment. Our findings underscore the critical role of Y RNAs on monocyte surfaces in capturing extracellular histones, thereby enhancing *IL-6* expression. Consistent with this model, targeted degradation of surface Y RNAs, facilitated by HASTE utilizing RNase H and DNA probes complementary to Y RNAs, led to a more pronounced inhibition, surpassing the efficacy of total surface RNA depletion for both *IL-6* (Figure 6D) and *IL-10* (Figure S14A) expression in human PBMC monocytes. Other surface RNAs beyond Y RNAs may play contrasting regulatory roles, potentially shielding cells from histone stimulation. Indiscriminate depletion of all surface RNAs could complicate effects. Future studies are required to understand roles of individual groups of surface RNAs in tuning cytokine expression.

### Extracellular histone supplementation triggers NF-κB and AP-1 activation

To decipher the signaling cascade triggered by extracellular histone stimulation, mRNA sequencing was performed on samples from THP-1 macrophages treated with either no stimuli (control group) or histone stimulation. KEGG (Kyoto Encyclopedia of Genes and Genomes) enrichment analysis identified key biological pathways, including cytokine, chemokine, NF-κB, TLR, RA, and MAPK signaling pathways (Figure 6E). Of particular interest are the NF-κB and MAPK pathways, as previous studies have reported NF-κB and Activator protein 1 (AP-1) activation in response to histone stimulation.^48^

To confirm these findings, THP-1 macrophages were pre-treated with the NF-κB inhibitor JSH-23, the AP-1 inhibitor SR 11302, or a combination of both inhibitors, prior to histone stimulation. As predicted, all three treatments significantly reduced the expression of *IL-6* gene in THP-1 macrophages induced by histone supplementation (Figure 6F), indicating the involvement of NF-κB and AP-1 in histone-triggered innate immune activation.

We propose a working model (Figure 7) in which surface Y RNAs on human monocytes facilitate the enrichment of extracellular histones released from nearby ruptured cells, thereby promoting IL-6 transcription and protein secretion through NF-κB and AP-1 activation via TLR2/4. Notably, targeted depletion of surface Y RNAs specifically inhibited IL-6 transcription and secretion (Figures 6A, C, D), with minimal effects on other cytokines such as GM-CSF, TNFα, and IL-1β (Figure S14B). This suggests the potential involvement of other unidentified factors, in the capture and presentation of extracellular histones.

**Figure 7.**
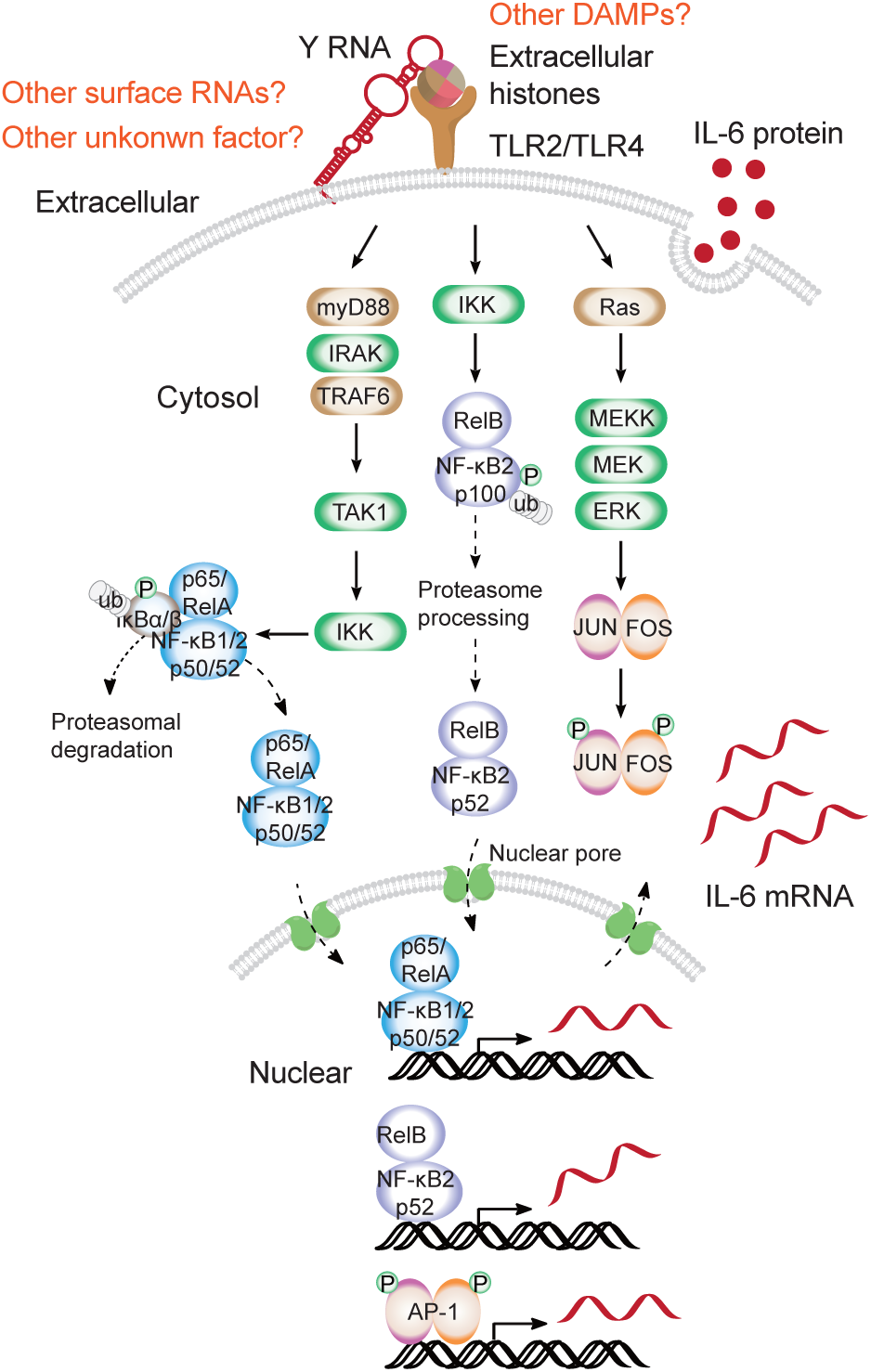
Working model depicting that surface Y RNAs enrich extracellular histones released from nearby ruptured cells, thereby promoting IL-6 gene transcription and protein secretion through NF-κB and AP-1 activation via TLR2/4.

## Discussion

In summary, our work significantly bolsters the evidence affirming the presence of surface-anchored RNAs on the outer membrane of primary cells. Our comprehensive portrayal of the surface RNA landscape within mammalian blood cells establishes a robust foundation for further investigations aiming at uncovering biological implications and functional significance of surface RNAs. Notably, our investigation unveils the residence of Y RNAs on surfaces of monocyte and B cells. The enrichment of extracellular histones through surface Y RNAs on human monocytes triggers NF-κB and AP-1 activation, leading to subsequent IL-6 expression via TLR2/4, revealing a previously unrecognized signaling pathway. Remarkably, targeted depletion of surface Y RNAs specifically inhibited IL-6 transcription and protein secretion (Figures 6A, C, D).

While significant progress has been made in validating and understanding surface RNAs, particularly the abundant Y RNAs displayed on the outer membrane of monocytes, many aspects of surface RNA biology across various cell types remain unexplored. For instance, Y RNAs are also significantly enriched on the surfaces of human primary B cells. Investigating the role of Y RNAs in adaptive immunity, including B cell activation and antibody secretion, will be intriguing directions in the future. The biological functions of other proteins interacting with surface Y RNAs, aside from histones, require further investigation. Several prevalent long non-coding surface RNAs, such as Mt-rRNA, may harbour equally significant functions warranting further investigation. Among these, U snRNAs (U1, U2, U4, U5, and U6 RNAs) integral components of U RNPs, are known as Sm antigens in the sera of SLE patients.^50^ Autoantibodies targeting aminoacyl-tRNA synthetases are implicated in the pathogenesis of anti-synthetase syndrome (ASS).^51^ Xist ribonucleoproteins have recently been reported to promote female sex-biased autoimmunity.^52^ Considering that surface RNAs are cell type-specific, methodologies capable of profiling both intracellular and surface RNAs simultaneously at the single-cell level are highly desirable.

Moving forward, we anticipate a diverse spectrum of roles for surface RNAs within distinct cell types, spanning neurons, germ cells, cancer cells, and beyond. Deciphering the intricate workings of surface RNAs necessitates continued investigation and exploration. Notably, the strategies developed in this study provide a foundational framework that will facilitate further advancements in this field.

### Limitations of the study

The methods developed in this study for profiling and imaging surface RNAs, as well as investigating surface RNA-binding proteins, employ live cells with a viability of no less than 95%. Consequently, live cells freshly dissected from tissue are required for surface RNA investigations. This limitation restricts the application of these techniques to frozen samples or formalin-fixed, paraffin-embedded (FFPE) samples, due to the risk of cell rupture and subsequent contamination from intracellular RNA. To address these limitations, it is crucial to develop additional methods that employ specialized strategies for investigating surface RNAs in frozen cells and FFPE samples. Furthermore, we investigated the biological function of surface Y RNA in human PBMCs in this study. Further exploration of surface Y RNAs in mouse models is highly desirable to determine whether the enrichment of extracellular DAMPs is conserved across human and murine systems.

## Resource availability

### Lead contact

Further information and requests for resources and reagents should be directed to and will be fulfilled by the Lead Contact, Lulu Hu.

### Materials availability

All plasmids generated in this study are available upon request.

### Data and code availability Data

The raw sequencing data generated from human blood cells reported in this paper have been deposited in the Genome Sequence Archive (Genomics, Proteomics & Bioinformatics 2021) in National Genomics Data Center (Nucleic Acids Res 2022), China National Center for Bioinformation / Beijing Institute of Genomics, Chinese Academy of Sciences (GSA-Human: HRA006257) that are publicly accessible at https://ngdc.cncb.ac.cn/gsa-human.

The sequencing data generated from mouse blood cells and cell lines referenced in this study have been meticulously archived and made publicly available via the Gene Expression Omnibus repository under the accession numbers GSE252429 (https://www.ncbi.nlm.nih.gov/geo/query/acc.cgi?acc=GSE252429, token: odypaokipjivzmv), GSE252682 (https://www.ncbi.nlm.nih.gov/geo/query/acc.cgi?acc=GSE252682, token: slqbscuunxkbbwt), and GSE262356 (https://www.ncbi.nlm.nih.gov/geo/query/acc.cgi?acc=GSE262356, token: kpkpayeivxkdbcr).

### Code

The original code has been deposited at GitHub: https://github.com/xuchu16/Surface-RNA. For inquiries regarding additional information needed for the reanalysis of the data presented in this paper, interested parties may obtain the requisite details from the lead contact upon request.

## Experimental model and subject details

### Mice

C57BL/6 mice were bred and housed in a specific pathogen–free environment within the animal facility. All animal care and experimental procedures adhered to guidelines and regulations approved by the Animal Care and Use Committee of Fudan University School of Medicine. Ethical principles and welfare standards were strictly maintained during the breeding, maintenance, and experimental manipulation of the mice.

### Subjects

Human cord blood samples were ethically collected from patients at the Obstetrics and Gynecology Hospital of Fudan University, with prior informed written consent. Collection procedures followed approved guidelines of the Ethics Committee of the Institutes of Biomedical Sciences, Fudan University, ensuring stringent ethical standards and privacy considerations. Human peripheral blood samples were ethically collected from healthy adults at the Traditional Chinese Medicine Hospital of Shijiazhuang, with prior informed written consent. The collection procedures adhered to the approved guidelines of the Clinical Trial Ethics Committee of the hospital.

## Acknowledgements

This work was supported by National Key R&D Program of China (2024YFA1307800, 2021YFA1100400), General Program of National Natural Science Foundation of China (32471340), Natural science foundation of Science and Technology Commission of Shanghai Municipality (STCSM, 21ZR1480300), and Fudan University Start-up funding to L.H. This work was also supported by General Program of National Natural Science Foundation of China (82070107), the National Excellent Youth Fund of National Natural Science Foundation of China (82222005), and Fudan University Start-up funding to X. H. We thank Dr. D.Y. and Dr. L.C. for advice in THP-1 cell culture and differentiation. We thank Dr. M.Y. from the Obstetrics and Gynecology Hospital of Fudan University for assisting with the collection of human umbilical cord blood samples. We thank Dr. H. M. for sharing pLKO plasmid. We thank The Professional Technical Service Platform for Critical Diseases in Shanghai, China (No.22142202400) for mass spectrometry analysis. We thank the Core Facility of Shanghai Medical College, Fudan University for providing the instruments used in this work.

## Contributions

X.J. performed all the experiments of method development, confocal imaging, surface RNA profiling, THP-1 cell line activation, analysed the data, and prepared figures. C.X. analysed the high-throughput sequencing data and prepared the figures. E.Y performed all the innate immune activation experiments using human monocytes purified from PBMC. D.X. prepared all types blood cells form human umbilical cord blood for AMOUR analysis, and Y RNA specific single cell sequencing under the guidance of X.H. Y.P. helped C.X. in data analysis under C. H’s guidance. J.S. helped X.J. in the real-time confocal imaging and BFA inhibition experiments. Z.L., Q.S., Q.C., and W.H. helped X.J in the confocal imaging and mass spectrometry experiments. X.H. helped X.J. in the single cell sequencing experiments under the guidance of Y.X. S.H. helped J.X. in the human histone protein preparation. Y.X. and C.H. proofread the manuscript. L.H. conceived and supervised the project, and wrote the manuscript with input from all authors.

## Ethics declarations

Human cord blood samples were ethically collected from patients at the Obstetrics and Gynecology Hospital of Fudan University, with prior informed written consent. Collection procedures followed approved guidelines of the Ethics Committee of the Institutes of Biomedical Sciences, Fudan University. Human peripheral blood samples were ethically collected from healthy adults at the Traditional Chinese Medicine Hospital of Shijiazhuang, with prior informed written consent. The collection procedures adhered to the approved guidelines of the Clinical Trial Ethics Committee of the hospital. All sample collections adhere to stringent ethical standards and privacy considerations.

## Supplemental information

**Figure S1 Localization of RNA molecules on the outer membrane surface of cell lines and primary human cells, identified by Intact-Surface-FISH, related to Figure 1**

(A) Schematic flow chart of Intact-Surface-FISH. Live primary cells or cell lines are incubated with Cy3-DNA probes complementary to target surface RNAs, washed, and subjected to flow cytometry analysis or confocal imaging.

(B) Transmission-through-dye confocal imaging of live HEK 293T cells under various treatments: control (upper panel), Intact-Surface-FISH (middle panel), and 0.2% Triton X-100 treatment prior to Intact-Surface-FISH for 5 minutes (bottom panel). Cells were stained with the membrane-permeable dye CellTracker Orange and the membrane-impermeant quencher acid blue 9 (AB9). In cells with intact membranes, AB9 is excluded, and CellTracker Orange fluorescence is observed throughout the cell. In cells with compromised membranes, AB9 enters and quenches CellTracker Orange, reducing or eliminating fluorescence signals. Box plots illustrate the mean fluorescence intensity per image for the control group, Intact-Surface-FISH-treated group, and 0.2% Triton X-100 permeabilized group.

(C) Confocal imaging of HeLa and HEK 293T cells stained with Cy3-Lambda DNA control and Cy3-N20 DNA probe (N represents a random A/T/C/G), without (middle panel) and with (bottom panel) RNase A & T1 pre-treatment, using the Intact-Surface-FISH strategy. Quantitative analysis of mean fluorescence intensity per cell is provided for each treatment. n indicates the number of cells examined.

(D) Fluorescence labeling of live hUCB-MNCs with Cy3-Lambda DNA control and Cy3-N20 DNA probes complementary to surface RNAs, using Intact-Surface-FISH. The PBS-incubated group serves as a negative control, while 4% PFA-fixed cells permeabilized with 0.2% Triton X-100 prior to Cy3-N20 incubation serve as the positive control group. The gated region indicates the cell population displaying specific RNAs on the cell surface.

(E) Mean fluorescence intensity (MFI) of live hUCB-MNCs stained with Cy3-Lambda DNA control and Cy3-N20, with or without Triton X-100 permeabilization, identified using Intact-Surface-FISH.

(F) Confocal imaging of hUCB-MNCs stained with Cy3-Lambda DNA or Cy3-N20, with or without Triton X-100 permeabilization, using Intact-Surface-FISH. Quantitative analysis of mean fluorescence intensity per cell is provided for each treatment. n indicates the number of cells examined. All flow cytometry analyses and confocal imaging data represent three independent experiments with similar results, shown as mean ± s.d., and analyzed using an unpaired two-tailed Student’s *t*-test; ns: not significant; **: p < 0.01; ***: p < 0.001; ****: p < 0.0001.

**Figure S2 Validation of the AMOUR strategy in HEK 293T cells using confocal imaging, related to Figure 1**

(A) Transmission-through-dye confocal imaging for HEK 293T cells under different treatments: live cell control group (upper panel), AMOUR-treated group (middle panel), and 0.2% Triton X-100 permeabilized group (bottom panel). Cells with intact membranes show bright fluorescence signals, while cells with damaged membranes exhibit quenched fluorescence signals (left panel). Box plots illustrate the mean fluorescence intensity per cell for the control group, AMOUR-treated group, and the group permeabilized with 0.2% Triton X-100 prior to AMOUR treatment (right panel).

(B) Confocal imaging of Alexa Fluor 647-labeled primary GAPDH antibody staining in HEK 293T cells under different treatments (left panel): AMOUR-treated group (upper panel) and 0.2% Triton X-100 permeabilized group (bottom panel). Box plots (right panel) display the mean fluorescence intensity per image for the AMOUR-treated group, and the group permeabilized with 0.2% Triton X-100 prior to AMOUR treatment.

(C) Confocal imaging of Cy3-labeled DNA probes complementary to GAPDH mRNA entry into HEK 293T cells under different treatments (left panel): AMOUR-treated group (upper panel), and the group permeabilized with 0.2% Triton X-100 prior to AMOUR treatment (bottom panel). Box plots (right panel) show the mean fluorescence intensity per image for the AMOUR-treated group, and the group permeabilized with 0.2% Triton X-100 prior to AMOUR treatment.

(D) Confocal imaging of Alexa Fluor 647-labeled T7 RNA polymerase entry into HEK 293T cells under different treatments (left panel): AMOUR-treated group and Triton X-100 permeabilized groups at concentrations of 0.001%, 0.01%, and 0.2% prior to AMOUR treatment. Box plots (right panel) depict the mean fluorescence intensity per image for the AMOUR-treated group, and the group permeabilized with Triton X-100 prior to AMOUR treatment.

The imaging experiments were repeated three times independently with similar results. Data are shown as mean ± s.d., and analyzed using an unpaired two-tailed Student’s *t*-test; ns: not significant; ****: p < 0.0001.

**Figure S3 Validation of the AMOUR strategy in HEK 293T cells using high-throughput sequencing, related to Figure 1**

(A) Reads coverage of a 352 nt RNA model amplified utilizing AMOUR strategy.

(B) Heatmap showing the correlation among specific RNAs identified by AMOUR with 3% HEK 293T cell lysate, AMOUR control, and RNase A & T1 treatment prior to AMOUR in HEK 293T cells (These datasets, generated using the AMOUR technique following various treatments, are intended specifically for method validation. They are distinct from the datasets derived from HEK293T cells presented in Figure 1 and Figure S4).

(C) Heatmap displaying specific RNAs identified by AMOUR with 3% HEK 293T cell lysate, AMOUR control, and RNase A & T1 treatment prior to AMOUR in HEK 293T cells.

(D) The predicted minimum folding free energy (MFE) is plotted for transcripts enriched following RNase A and RNase T1 treatment prior to AMOUR in HEK 293T cells, AMOUR control, and 3% HEK 293T cell lysate groups. The number of transcripts in each group is provided at the bottom.

(E) Box plots depict the GC content of transcripts enriched following RNase A and RNase T1 treatment prior to AMOUR in HEK 293T cells, AMOUR control, and 3% HEK 293T cell lysate groups. The number of transcripts in each group is provided at the bottom.

(F) Box plots display TPM (transcripts per million) expression values across various RNA categories in the AMOUR control and 3% HEK 293T cell lysate groups. Transcripts with a TPM value greater than 50 in at least one biological sample were included in the analysis. The number of transcripts in each RNA category is shown at the bottom.

(G) Box plots showing TPM expression values of representative transcripts identified in the AMOUR control, 3% HEK 293T cell lysate, and RNase A and RNase T1 treatment prior to AMOUR groups.

Data are presented as the mean ± standard deviation (s.d.) and were analyzed using an unpaired two-tailed Student’s *t*-test; ns, not significant; *: p < 0.05, t; ****, p < 0.0001.

**Figure S4 Characteristics of surface RNAs on the outer membrane of HEK 293T and Hela cells, related to Figure 1**

(A) Pearson correlation analysis of two independent replicates of AMOUR datasets for cultured HEK 293T cells, excluding tRNA (These datasets in Figure S4, generated from HEK 293T cells, are consistent with those presented in Figure 1 but distinct from those shown in Figure S3.).

(B) Pie chart showing the proportions of transcript types (only those with TPM > 50) identified using AMOUR in HEK 293T cells, including tRNA.

(C) Bar plot showing transcript type statistics of surface RNAs identified using the AMOUR strategy (only those with TPM > 50) for HEK 293T cells, excluding tRNA.

(D) Bar plot showing transcript counts of representative HEK 293T surface RNA molecules.

(E) Venn plot demonstrating the overlap between surface tRNA detected via AMOUR and glycosylated tRNA reported by Ryan A Flynn et al. (Cell. 2021).

(F) Scatter plot highlighting the overlap between surface tRNAs detected via AMOUR and glycosylated tRNA reported by Ryan A Flynn et al. (Cell. 2021). AMOUR-specific surface tRNAs are indicated in blue, whereas common tRNAs are shown in orange.

(G) Heatmap depicting the correlation of surface RNAs identified in HEK 293T and HeLa cells using AMOUR.

(H) Pie chart showing the proportions of transcript types (only those with TPM > 50) identified using AMOUR in Hela cells, including tRNA.

(I) Box plots display TPM expression values across various RNA categories in the HEK 293T and HeLa cells identified with AMOUR. Transcripts with a TPM value greater than 50 in at least one biological sample were included in the analysis. The number of transcripts in each RNA category is shown at the bottom.

Data are presented as the mean ± standard deviation (s.d.) and were analyzed using an unpaired two-tailed Student’s *t*-test; ns, not significant; *, p < 0.05; ****, p < 0.0001.

**Figure S5 Validation of ARMOUR via comparative profiling of surface RNAs using biotin-WGA pull down, related to Figure 1**

(A) Schematic flowchart illustrating the Wheat Germ Agglutinin (WGA) assisted membrane-associated RNA pull-down (WGA-Pd) strategy. Biotin-WGA is utilized to enrich membrane-associated RNA, followed by library construction and sequencing.

(B) Heatmap showcasing the correlation among surface RNAs, excluding tRNA, identified by the AMOUR and WGA-Pd strategies for HEK 293T cells.

(C) Heatmap displaying the correlation among surface tRNAs identified by the AMOUR and WGA-Pd strategies for HEK 293T cells.

(D) Pie chart showing the proportions of transcript types (only those with TPM > 50) identified by the AMOUR and WGA-Pd strategies for HEK 293T cells, excluding tRNA.

(E) IGV tracks displaying the reads coverage of representative surface RNAs: RNY1, RNY3, RNY4, and RNY5 identified by the AMOUR and WGA-Pd strategies for HEK 293T cells.

**Figure S6 Validation of surface RNA localization using Intact-Surface-FISH, related to Figure 2**

(A) Fluorescence co-labeling of live or fixed and permeabilized hUCB-MNCs with Cy5-DNA probes complementary to mRNA GAPDH and Cy3-DNA probes complementary to RNA Y5, using Intact-Surface-FISH. The gated region indicates the cell population displaying positive Cy3 and Cy5 signals. Quantitative analysis of the mean fluorescence intensity within the Cy3/Cy5-positive region is provided for each treatment. Data represent four independent experiments.

(B) Confocal co-imaging of live or fixed and permeabilized hUCB-MNCs with Cy5-DNA probes complementary to mRNA GAPDH and Cy3-DNA probes complementary to RNA Y5, using Intact-Surface-FISH or conventional RNA-FISH. Quantitative analysis of the fluorescence intensity per image is provided for each treatment.

(C) Confocal co-imaging of fixed and permeabilized hUCB-MNCs using Cy3-DNA probes complementary to representative RNAs, performed with conventional RNA-FISH. Quantitative analysis of the fluorescence intensity per image is provided for each treatment. All imaging experiments were conducted independently in triplicate, with similar results observed across all repetitions.

**Figure S7 Analysis of surface RNA transport via vesicle trafficking inhibition, related to Figure 2**

(A) Fluorescence-activated cell sorting (FACS) analysis of untreated Raji cells, as well as Raji cells pre-treated with Brefeldin A or RNase A & T1, followed by staining with the J2 dsRNA antibody. The left panel displays the FACS results, with the dashed vertical line indicating the J2-high population. Quantitative analysis of the percentage of cell fractions (middle panel) and mean fluorescence intensity (right panel) within the J2-high region is provided for each treatment condition.

(B) Confocal imaging of Raji cells pre-treated with Brefeldin A or RNase A & T1, followed by staining with the J2 dsRNA antibody. The left panel shows representative imaging results, while the right panel presents quantitative analysis of fluorescence intensity per image for each treatment condition.

All imaging and flow cytometry experiments were conducted independently in triplicate, with similar results observed across all repetitions. Data are presented as the mean ± standard deviation (s.d.) from three independent experiments and were analyzed using an unpaired two-tailed Student’s *t*-test; ***: p < 0.001; ****: p < 0.0001.

**Figure S8 Statistics of surface RNAs across diverse blood cell types, related to Figure 3**

(A-B) Stacked plots illustrating the distribution of RNA types among distinct human (A) and murine (B) blood cell subsets, excluding tRNAs.

(C) Euclidean distance analysis representing the dissimilarities in surface RNA datasets across distinct blood cell types, excluding tRNAs.

(D-E) Box plots presenting distinct highly abundant surface RNA profiles characterizing individual human (D) and murine (E) blood cell populations.

(F) Box plot showcasing prevailing highly abundant surface RNA molecules collectively observed across all analysed blood cell types in both human and mouse.

(G-I) Heatmap illustrating surface tRNAs identified within distinct human (G) and murine (H) blood cell subsets, along with those conserved across all analysed cell types in both human and mouse (I). Highlighted representative tRNA signatures emphasize discernible expression patterns within these subsets.

(J) Box plots depict TPM (transcripts per million) expression values across various RNA categories in B cells, monocytes, and T cells, as identified through total RNA sequencing. Transcripts with an average TPM value greater than 50 were included in the analysis. The number of transcripts in each RNA category is shown at the bottom.

Data are presented as the mean ± standard deviation (s.d.) and were analyzed using an unpaired two-tailed Student’s *t*-test; ns, not significant; *: p < 0.05.

**Figure S9 Pearson correlation analysis of three independent biological replicates of AMOUR datasets across human blood cell types, related to Figure 3**

**Figure S10 Pearson correlation analysis of three independent biological replicates of AMOUR datasets across murine blood cell types, related to Figure 3**

**Figure S11 Y RNAs capture histones, related to Figure 5**

(A) Schematic diagram of the <u>s</u>urface <u>R</u>NA interacting protein <u>c</u>apture assay (SRIC).

(B) Schematic diagram of the modified <u>R</u>NA interacting protein <u>c</u>apture assay (modified-RIC).

(C) Scatter plot illustrating proteins associated with the surface Y RNA mix (Y1/3/4/5). The dashed line indicates a fold change (+UV/-UV) threshold of 1.5. Proteins with significant mass spectrometry signal in the experimental group (+UV) are highlighted in orange. Data are derived from two independent biological replicates.

(D) Scatter plot showing surface Y1, Y3, and Y4 RNA associated proteins identified using modified RIC strategy. The dashed line represents the foldchange(+UV/-UV) of peptide fingerprint signal = 2. Histones, Ro60, and SSB (La) proteins are highlighted in orange. The data were analyzed based on two independent biological replicates.

(E) Saturation plot illustrating the binding of 3’FTSC-Y5 RNA with H2A-H2B and H3-H4 complex.

**Figure S12 HASTE effectively eliminate Y RNAs on the monocyte surface, related to Figure 6**

(A) PAGE illustrating enzyme efficacy towards 100 ng of Y4 RNA under varying conditions: lane 1: RNase H with c-Y4 RNA probe in 1 × commercial RNase H buffer; lane 2: RNase H with c-Y RNA probe in 1× PBS with 4 mM MgCl_2_; lane 3: RNase H with c-Y RNA probe in 1 × PBS with 1 mM MgCl_2_; lane 4: RNase H with c-Y RNA probe in 1 × PBS with 1% BSA and 4 mM MgCl_2_; lane 5: RNase H with c-Y RNA probe in 1 × PBS with 1% BSA and 1 mM MgCl_2_; lane 6: RNase A and RNase T1 in 1 × PBS; lane 7: RNase A and RNase T1 in 1 × PBS with 1% BSA; lane 8: c-Y4 RNA DNA probe; lane 9: Y4 RNA control without any treatment.

(B) Confocal imaging of THP-1 monocytes stained with Cy3-Lambda DNA or Cy3-DNA probes complementary to the Y RNA mix (Y1/3/4/5), with or without HASTE pre-treatment, using Intact-Surface-FISH. White arrowheads indicate specific locations where Intact-Surface-FISH signals of the Y RNA mix colocalize with the plasma membrane. Quantitative analysis of mean fluorescence intensity per cell is provided for each treatment.

(C) Fluorescence labeling of live hUCB-MNCs with Cy3-Lambda DNA control or Cy3-DNA probes complementary to the Y RNA mix, with or without HASTE pre-treatment, using Intact-Surface-FISH. The gated region indicates the cell population displaying Y RNAs on the cell surface. Quantitative analysis of the percentage of cell fractions within the Cy3-positive region is provided for each treatment.

(D) Confocal imaging of hUCB-MNCs stained with Cy3-Lambda DNA control or Cy3-DNA probes complementary to the Y RNA mix, with or without HASTE pre-treatment, using Intact-Surface-FISH. White arrowheads indicate specific locations of Intact-Surface-FISH signals. Quantitative analysis of mean fluorescence intensity per image is provided for each treatment.

All imaging experiments were performed independently in triplicate, with consistent results observed across all repetitions. Flow cytometry experiments were conducted independently with four replicates. Data are presented as the mean ± standard deviation (s.d.) and were analyzed using an unpaired two-tailed Student’s t-test; ns: not significant, **: p < 0.01, ***: p < 0.001, ****: p < 0.0001.

**Figure S13 Surace RNA depletion is non-disruptive, related to Figure 6**

(A-B) Brightfield images (A) and corresponding quantitative analysis (B) depicting the cell number and size of THP-1 monocytes: control group (left panel), experimental groups post total surface RNA depletion (middle panel), and Y RNA depletion (right panel).

(C) Cell viability analysis of the THP-1 monocytes: control group, experimental groups post total surface RNA depletion, and Y RNA depletion.

(D) mRNA levels of macrophage marker genes CCR5 (left panel), CD14 (middle panel), and CD36 (right panel) were determined via RT-qPCR in THP-1 macrophages differentiated from THP-1 monocytes with 100 ng mL^-1^ PMA for 48 hours. The groups analyzed include control, total surface RNA depleted, and Y RNA depleted (normalized to untreated control, n = 3).

(E) mRNA levels of GAPDH, IL-6 and IL-10 in THP-1 macrophages were measured after 6 hours of culture without calf thymus histone supplementation, following various pre-treatments, as determined by RT-qPCR. NC: untreated control group; RNase A & T1: THP-1 macrophages treated with RNase A and T1 for 30 minutes; RNase H & probes: THP-1 macrophages treated with RNase H and Y RNA complementary DNA probes for 30 minutes. Data are normalized to the NC group, n = 3. ND: not detachable.

All confocal imaging data represent three independent experiments with similar results, shown as mean ± s.d., and analyzed using an unpaired two-tailed Student’s *t*-test; ns: not significant, *: p < 0.05, ****: p < 0.0001.

**Figure S14 Surface Y RNAs on human monocytes facilitate IL-10 transcription, but not GM-CSF, TNFα, or IL-1β, in response to histone stimulation, related to Figure 6**

(A) mRNA levels of IL-10 in human monocytes purified from four human PBMC samples after 6 hours of supplementation with 50 μg mL^-1^ calf thymus histones, subsequent to various pre-treatments, determined by RT-qPCR. NC: untreated control group; PC: human monocytes treated with histones without any pre-treatment; RNase A & T1: human monocytes treated with RNase A and T1 for 30 minutes prior to histones supplementation; HASTE: human monocytes treated with RNase H and surface Y RNA complementary DNA probes for 30 minutes prior to histone supplementation. Data normalized to the NC group, n = 3.

(B) mRNA levels of GM-CSF, TNFα, and IL-1β in THP-1 macrophages after 6 hours of supplementation with 50 μg mL^-1^ calf thymus histones, subsequent to various pre-treatments, determined by RT-qPCR. NC: untreated control group; PC: THP-1 macrophages treated with histones without any pre-treatment; RNase A & T1: THP-1 macrophages treated with RNase A and T1 for 30 minutes prior to histones supplementation; HASTE: THP-1 macrophages were treated with RNase H and DNA probes complementary to surface Y RNAs for 30 minutes prior to histone supplementation. Data normalized to the NC group, n = 3.

All data represent three independent experiments, shown as mean ± s.d., and analyzed using an unpaired two-tailed Student’s *t*-test; ns: not significant; **: p < 0.01; ***: p < 0.001; ****: p < 0.0001.

